# GTPase activity-coupled treadmilling of the bacterial tubulin FtsZ organizes septal cell-wall synthesis

**DOI:** 10.1101/077610

**Authors:** Xinxing Yang, Zhixin Lyu, Amanda Miguel, Ryan McQuillen, Kerwyn Casey Huang, Jie Xiao

## Abstract

The bacterial tubulin FtsZ is the central component of the division machinery, coordinating an ensemble of proteins involved in septal cell-wall synthesis to ensure successful constriction. How cells achieve this coordination is unknown. We used a combination of imaging, genetic and biochemical approaches to demonstrate that in *Escherichia coli* cells FtsZ exhibits dynamic treadmilling predominantly determined by its GTPase activity, and that the treadmilling dynamics directs processive movement of the septal cell-wall synthesis machinery. In FtsZ mutants with severely reduced treadmilling, the spatial distribution of septal synthesis and the molecular composition and ultrastructure of the septal cell wall are substantially altered. Thus, the treadmilling of FtsZ provides a novel and robust mechanism for achieving uniform septal cell-wall synthesis to enable correct new pole morphology.

**One-sentence summary:** The bacterial tubulin FtsZ uses GTP hydrolysis to power treadmilling, driving processive synthesis of the septal cell wall.

## Main Text

The tubulin homolog FtsZ (1) is the central component of the cell division machinery in nearly all walled bacterial species (2). During division, FtsZ polymerizes on the cytoplasmic face of the inner membrane to form a ring-like structure (the Z-ring) and recruits more than thirty proteins to the division site, many of them involved in septal synthesis of the peptidoglycan (PG) cell wall (3). The GTPase activity of FtsZ is highly conserved, and the binding and hydrolysis of GTP underlie the dynamic assembly and disassembly of FtsZ polymers (4, 5). Hydrolysis dynamics also form the basis for several prevalent mechanisms by which the Z-ring could generate constriction forces (6). However, in *Escherichia coli* cells, GTPase activity of FtsZ appears nonessential for cell division: G105S (FtsZ84) cells constrict at the same rate as wildtype (WT) (7) despite an ~80-90% reduction in GTPase activity *in vitro* (8, 9), and a number of other GTPase mutants are viable despite various division defects (10). Thus, while it has been clearly demonstrated that FtsZ’s GTPase activity is coupled to the highly dynamic structural organization of the Z-ring (4, 11), the biological function of the conserved GTPase activity of FtsZ in bacterial cell division remains elusive. Here, we use a combination of imaging, genetic, and biochemical approaches to demonstrate that FtsZ’s GTPase activity powers treadmilling of the Z-ring, which directs the processive movement of the septal cell-wall synthesis machinery to ensure uniformly distributed incorporation of new cell wall, and hence plays a critical role in polar morphogenesis during cell division.

To understand the dynamic structural reorganization of the Z-ring, we first characterized WT Z-ring dynamics in live *E. coli* BW25113 cells by monitoring the fluorescence of FtsZ-GFP expressed in the presence of endogenous, unlabeled FtsZ (FtsZ-GFP is 46% ±4.3% of total FtsZ concentration, mean ± standard deviation, (µ ± s.d.), *n* = 3, fig. S1) using total internal reflection microscopy (TIRF) (12). The TIRF illumination confines the imaging window to a thin layer of ~80 nm in depth and ~500 nm in width at the bottom of the cell (fig. S2) (12), enabling high sensitivity and minimizing photobleaching. Interestingly, in contrast to the roughly constant fluorescence intensity of Z-rings commonly observed in wide-field fluorescence imaging, we found that the integrated TIRF intensity of the Z-ring exhibited large, approximately periodic fluctuations (Fig. 1A and B, fig. S3, Movies S1, S2). We observed similar behaviors using a different fluorescent protein fusion to FtsZ (fig. S4A, D) and of a GFP-ZapA fusion (fig. S4B) that binds to FtsZ and complements a null *zapA* mutant in the absence of labeled FtsZ (13). Furthermore, this behavior was absent in fixed cells (fig. S4C). Thus, combined with our previous observation that FtsZ-GFP and FtsZ co-assemble into polymers within the Z-ring (7), we conclude that the fluctuations in FtsZ-GFP fluorescence intensity represent overall FtsZ dynamics, and likely reflect repeated assembly and disassembly cycles of FtsZ polymers in the Z-ring.

**Figure 1:**
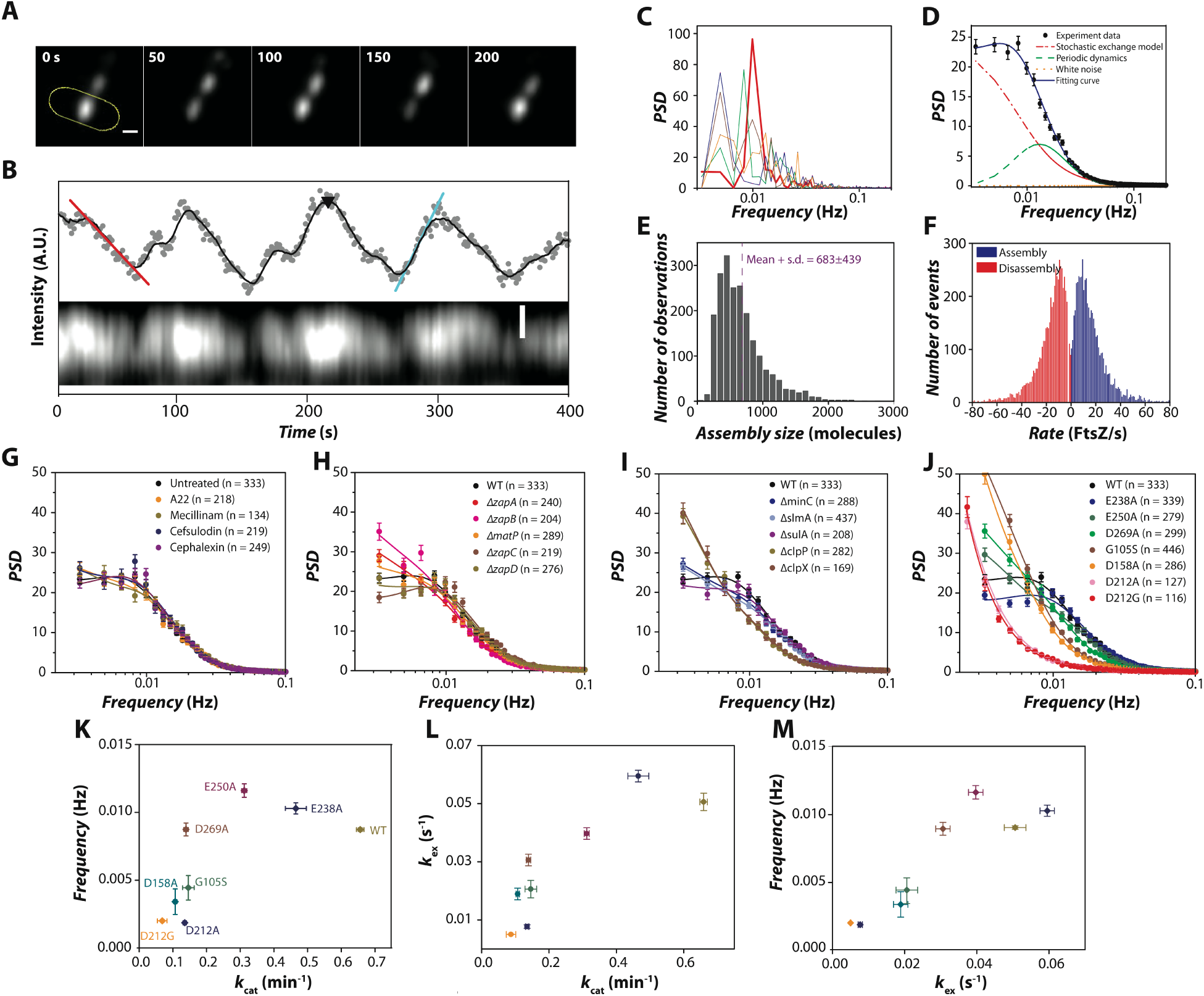
FtsZ polymers in the Z-ring exhibit periodic assembly and disassembly dynamics that are coupled to GTPase activity. (A) Top: Montages of live *E. coli* cells showing periodic FtsZ-GFP intensity fluctuations. The time-lapse movie (frame rate: 1/s, Movie S1) of the yellow-outlined cell at the bottom was used to generate the kymograph of Z-ring intensity in the TIRF imaging window and the integrated Z-ring fluorescence time trace (B). The black curve in (B) is the moving average (every 20 points) of the raw intensity (gray dots). The assembly and disassembly rates are determined as the maximal slopes along each rise (blue line) and decay (red), respectively. Assembly size, the maximum number of FtsZ molecules (FtsZ and FtsZ-GFP) in the imaging window during each period indicated by the black arrow head in (B), was estimated from the main intensity peaks using calibrations of FtsZ-GFP fluorescence and expression level (12). Scale bars: 0.5 µ m. Corresponding time-lapse movie and that of an additional cell were shown in Movies S1 and S2. (C) Representative power spectrum density (PSD) curves for individual cells, with one cell highlighted in solid red. (D) The mean PSD over all cells (black dots, error bars: standard error (s.e.), *n* = 333 cells), fitted with a model (blue curve) that takes into account stochastic subunit exchange between the Z-ring and the cytoplasmic pool (dashed red curve) and the periodic fluctuations (dashed green curve) (12). (D) Distribution of assembly size of FtsZ polymers. The mean is 683 ± 439 (s.d., n = 2039 fluorescence peaks), including both FtsZ and FtsZ-GFP. (E) Distributions of assembly (blue) and disassembly (red) rates. (G-J) Average PSD curves in drug-treated cells (G), in cells lacking Z-ring stabilizers (H) or regulators (I), and in cells expressing FtsZ GTPase mutants (J). Error bars: s.e. Sample size numbers (n) are indicated in the corresponding graphs. (K) Correlation between the GTPase catalytic turnover rate *k*_cat_ and the periodic frequency of FtsZ GTPase mutants. (L) Correlation between *k*_cat_ and the stochastic exchange rate *k*_ex_ of FtsZ GTPase mutants. (M) Correlation between *k*_ex_ and the periodic frequency of FtsZ GTPase mutants. Error bars: s.d.

We used the power spectrum density (PSD), a way to characterize the amplitude of different frequency components in a signal, to analyze the intensity fluctuations. We found that the PSDs of individual cells displayed clear peaks between 0.005-0.02 Hz (Fig. 1C), signifying characteristic periods of 50-200 s. The mean PSD across all cells (Fig. 1D, black dots; *n* = 333 cells) is significantly different from the monotonic decay (Fig. 1D, red dashed curve) predicted from a kinetic model in which only the stochastic exchange of FtsZ subunits between the Z-ring and the cytoplasmic pool is taken into account (4, 5, 12), indicating the presence of other dynamic processes of the Z-ring. Subtracting the contribution of stochastic subunit exchange from the mean PSD curve (12) resulted in a log-normal-like distribution (Fig. 1D, green curve) with a peak frequency of 0.0087 ± 0.0007 Hz (µ ± s.e. *n* = 333 cells, Table S1), corresponding to a period of 115 ± 10 s. Autocorrelation analysis of the time traces further confirmed the same time scale (fig. S5). To estimate the assembly size, which we define as the number of FtsZ molecules assembled in each fluorescence intensity peak in the TIRF imaging window (Fig. 1B, black arrow head), we calibrated the fluorescence intensity of FtsZ-GFP with cellular expression levels of FtsZ and FtsZ-GFP (fig. S1, S6) (12), and arrived at 683 ± 439 molecules (Fig. 1E, µ ± s.d., *n* = 2039 peaks, including both FtsZ and FtsZ-GFP). The large number of molecules suggests the presence of multiple FtsZ protofilaments, likely corresponding to the smaller FtsZ clusters previously observed via superresolution imaging (7). Notably, the assembly and disassembly rates of these FtsZ polymers estimated from the fluorescence rises (blue line in Fig. 1B) and falls (red line in Fig. 1B) in the periodic fluctuations, respectively, are essentially the same (assembly rate = 16.7 ± 14.6 FtsZ/s, µ ± s.d., *n* = 5393 fluorescence rises; disassembly rate = 16.8 ± 14.0 FtsZ/s, *n* = 5549 fluorescence falls; Fig. 1F), suggesting that the continuous assembly and disassembly of FtsZ polymers are in a dynamic steady state.

Previous studies have shown that the dynamic circumferential movement of MreB, an actin homolog that spatially organizes cell-wall synthesis (14), is driven by PG-synthesis enzymes in both *E. coli* (15) and *Bacillus subtilis* (16, 17). To test whether our observed dynamics of FtsZ polymers is also driven by PG synthesis, we treated cells with specific inhibitors of MreB (A22), the bifunctional division-specific glycosyltransferase and transpeptidase PBP1b (cefsulodin), the elongation-specific transpeptidase PBP2 (mecillinam), and the division-specific transpeptidase FtsI (PBP3) (cephalexin) at concentrations above their minimum inhibition concentration (12). In contrast to MreB, periodic FtsZ fluctuations quantified by PSD analysis remained identical to those in WT cells for all treatments (Fig. 1G, Table S2). To test whether the dynamics is influenced by other proteins that affect Z-ring assembly, we further examined ten mutant backgrounds individually lacking one of the proteins that regulates (SlmA, SulA, MinC, ClpX, and ClpP) or stabilizes (ZapA, ZapB, ZapC, ZapD and MatP) the Z-ring (12, 18). None of these genetic perturbations substantially affected Z-ring behavior (Fig. 1H,I, Table S3). These data suggest that the observed dynamics of the Z-ring are only marginally impacted by cell-wall synthesis or protein regulators, and instead are likely due to FtsZ’s intrinsic polymerization properties, which are known to be related to GTPase activity.

To examine whether GTPase activity influences the periodic assembly and disassembly dynamics, we constructed seven strains each containing a single point mutation at the chromosomal *ftsZ* locus (E238A, E250A, D269A, G105S, D158A, D212A, or D212G) (12). These mutants have previously been reported to alter GTP hydrolysis activity to different degrees (19). Using purified proteins, we measured these mutants’ GTPase activity *in vitro* under our experimental conditions (12, 20), and found that that their catalytic turnover rates (*k*_cat_) ranged from 14% to 71% of FtsZ^WT^ (fig. S7A, Table S1) (12). Note that because of the high cellular concentration of GTP (~5 mM) (21), the *in vivo* GTPase activity of these mutants should mainly represent their maximal GTP hydrolysis rate (reflected in *k*_cat_), even though there are differences in GTP-binding affinity (fig. S7A, Table S1).

Consistent with reports that defects in GTP hydrolysis severely impact FtsZ assembly *in vitro* (22), we found that the periodic assembly and disassembly dynamics of the Z-ring were significantly reduced in mutants with lower GTPase activity, and essentially abolished in the D212A and D212G mutants, which has the most severe reductions in activity (Fig. 1J, K, Table S1). In addition, the subunit-exchange rates (*k*_ex_) of these mutants extracted from the PSD curves (Fig. 1K) or using fluorescence recovery after photobleaching (FRAP) (fig. S7C to E) decreased with *k*cat (12), consistent with the expectation that these dynamics are coupled to GTP hydrolysis. We observed the same trend using GFP-ZapA as a marker for the unlabeled Z-ring in these mutants (fig. S4E). The fluctuation frequency and subunit exchange rate of each mutant were highly correlated with *k*_cat_ (*R*_*Spearman*_ = 0.81, *p* = 0.02 and *R*_*Spearman*_ = 0.93, *p* = 0.002, Fig. 1K, L, respectively), and with each other (*R*_*Spearman*_ = 0.90, *p* = 0.005, Fig. 1M). Clearly, the periodic dynamics of FtsZ polymers in the Z-ring is strongly coupled to GTP hydrolysis.

What type of assembly/disassembly process gives rise to the observed periodic behavior of FtsZ polymers? In some kymographs of cells lacking well-defined midcell Z-rings, zigzags of FtsZ-GFP fluorescence were readily visible, indicating directional FtsZ polymer movement (fig. S8A, B and Movies S3-6). By imaging at higher temporal and spatial resolution (12), we identified that in many cells FtsZ polymers exhibited transverse, processive movement across the short axis of the cell (Fig. 2A, B, C, fig. S9, Movies S6-11). This processive movement was particularly prominent in shorter cells (64% of cells with length < 2.8 µm, *n* = 53 cells, Fig. 2A, C), prior to the establishment of a stable midcell Z-ring. In longer cells (≥ 2.8 µm) in which the Z-ring is stably assembled at midcell, a smaller percentage of cells exhibited such dynamics (34%, *n* = 41 cells, Fig. 2B, C), likely because overlapping FtsZ polymers in the Z-ring could not be resolved under diffraction-limited imaging.

**Figure 2:**
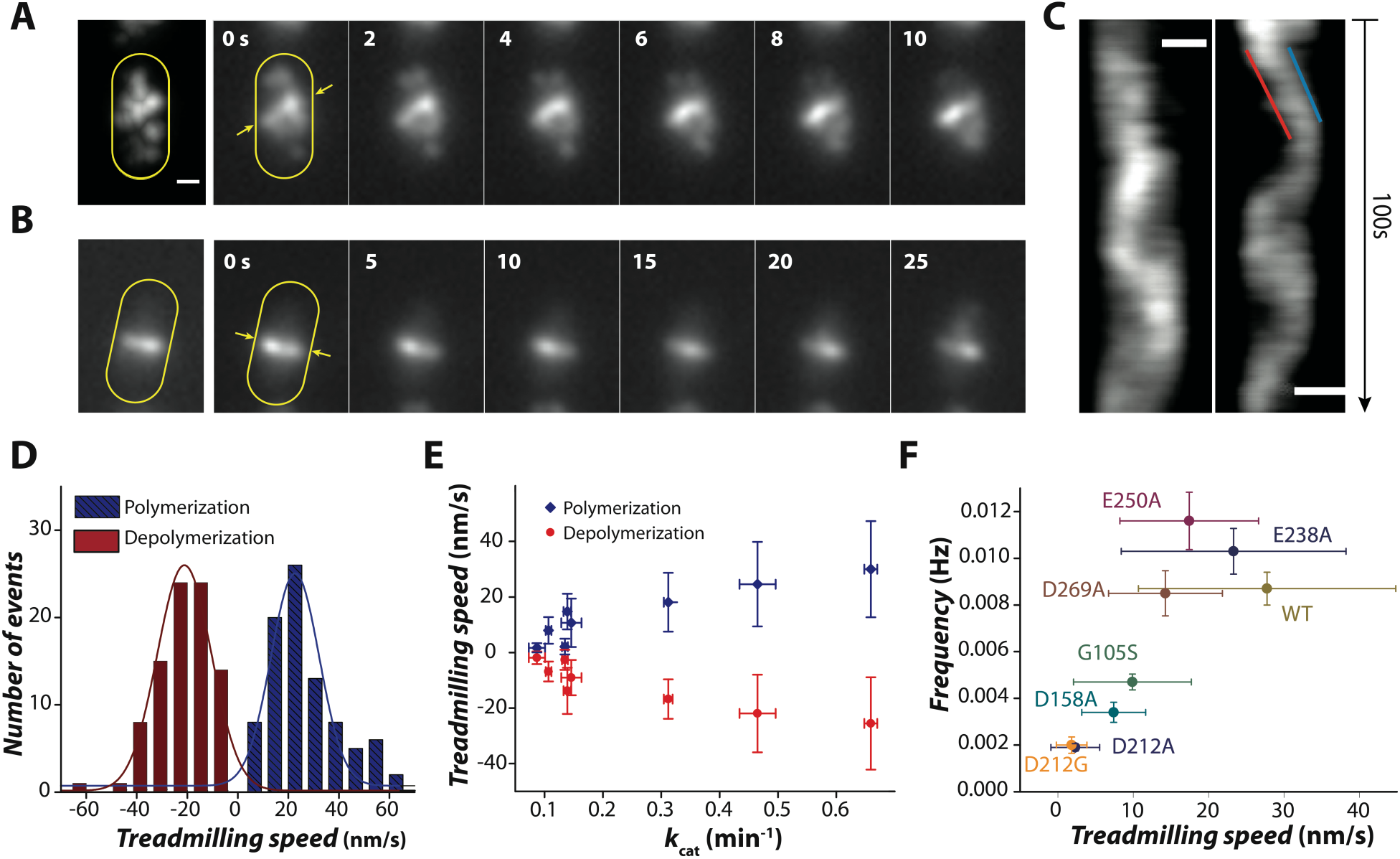
FtsZ polymers exhibit treadmilling dynamics in live *E. coli* cells. (A and B) Maximum intensity projection (left) and montages of the time-lapse movie (right, frame rate: 2/s, Movies S6, S7) of a cell in which a midcell Z-ring has not assembled (A) and another cell in which a clear midcell Z-ring is visible (B). The corresponding kymograph of each cell computed from the intensity along the line between the two yellow arrows in the first frame of the montage in (A) and (B) was shown in (C). The polymerization and depolymerization speeds were measured from the leading (blue line) and trailing (red) edges of individual cells’ kymographs as shown in (C), and the corresponding distributions were shown in (D). Scale bars: 0.5 µ m. (E) Polymerization and depolymerization speeds of FtsZ GTPase mutants correlate with *k*_cat_. Error bars: s.d. (F) Absolute treadmilling speed (average of both polymerization and depolymerization speeds) correlates with the periodic assembly/disassembly frequency of FtsZ polymers in FtsZ GTPase mutants. Error bars: s.d. (treadmilling speed) or s.e.(frequency).

We and other groups previously showed that individual FtsZ molecules remain stationary in assembled FtsZ polymers ((23–25)). Therefore, the observed processive movement of FtsZ polymers is most consistent with polymerization at one end and depolymerization at the other, an essential feature of treadmilling. Indeed, treadmilling of FtsZ polymers assembled on supported lipid bilayers *in vitro* has recently been observed (24). Using kymographs, we measured the apparent polymerization and depolymerization speeds of each trajectory (Fig. 2C) (12). We found that while there were substantial variations in the speeds among individual cells, the distributions of polymerization and depolymerization speeds across the population of cells were similar (Kolmogorov-Smirnov test, *p* = 0.073), with mean speeds of 30 ± 17 nm/s (µ ± s.d., *n* = 91 traces) and 26 ± 17 nm/s, respectively (Fig. 2D, Table S1). This behavior is consistent with the classic definition of treadmilling dynamics, and hereafter we combined these two speeds together and used the average as the treadmilling speed (Table S1). Importantly, we found that the treadmilling speed of each mutant is strongly correlated with the corresponding *k*_cat_ (*R*_*Spearman*_ = 0.95, *p* = 0.001, Fig. 2E), and with the frequency of FtsZ-GFP periodic fluctuations (*R*_*Spearman*_ = 0.88, *p* = 0.007, Fig. 2F). Thus, these data strongly imply that FtsZ’s GTPase activity underlies the treadmilling behavior, which is responsible for the periodic assembly and disassembly dynamics of FtsZ polymers.

Next, we investigated the role of FtsZ treadmilling in *E. coli* cell division. Using scanning electron microscopy (SEM), we noticed that FtsZ mutants with severe disruptions to GTPase activity exhibited abnormal septum morphology (Fig. 3A, fig. S10). In contrast to the smooth, symmetrically invaginated hemispherical septa in WT cells, mutant strains frequently exhibited slanted, twisted, and/or incomplete septa along the cell length and at the cell pole (Fig. 3A, fig. S10). It has previously been suggested that the pattern of FtsZ localization dictates the shape of the invaginating septum, based on the twisted and asymmetric septa observed in a temperature-sensitive FtsZ mutant (FtsZ26) that forms arcs and spirals rather than a ring along the membrane (26). Superresolution imaging in several bacterial species has revealed that the Z-ring is actually a discontinuous collection of FtsZ clusters (7, 11, 27–30), making it unclear how a smooth, hemispherical septum morphology is achieved. We hypothesize that treadmilling allows FtsZ polymers to evenly sample the surface of the growing septum over time, thereby ensuring a uniform spatial distribution of PG synthesis along the septum. In this scenario, Z-rings formed by arcs or clusters of GTPase mutants that essentially lack treadmilling ability, such as D212A/G, would not distribute septal PG synthesis evenly on average during constriction, resulting in incomplete and/or asymmetric septa.

**Figure 3:**
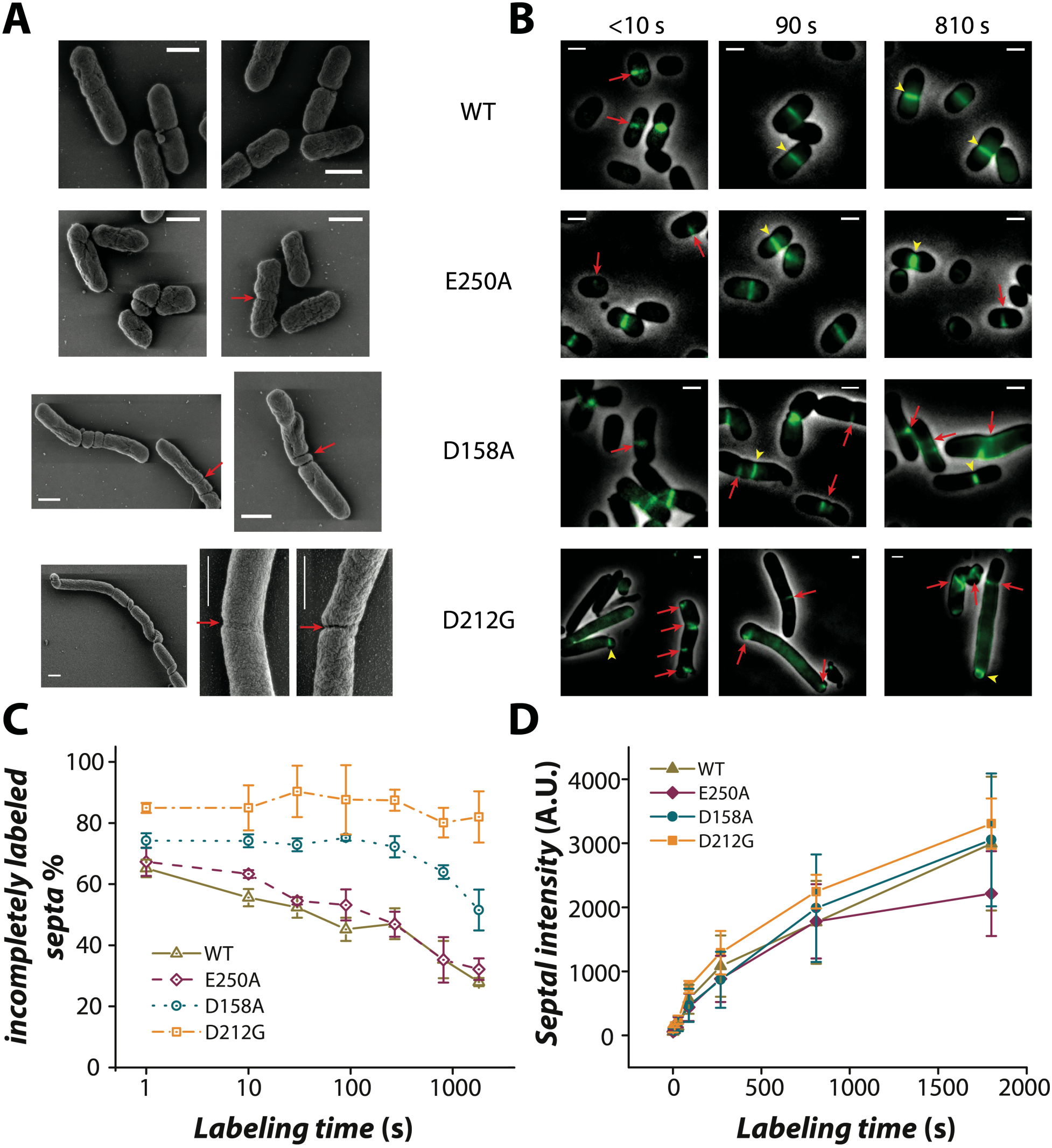
FtsZ GTPase mutants change the spatial distribution pattern but not the total amount of septal PG synthesis. (A) Representative SEM images of WT, E250A, D158A, and D212G cells. Arrows denote deformed, asymmetric septa. (B) Representative images of HADA-labeled septa for short (< 10 s), intermediate (90 s), and long (810 s) labeling pulses. Arrows and arrowheads denote incomplete and complete septa, respectively. (C) Severe GTPase mutants have large percentage of cells with incompletely labeled septa even for long pulses. Error bars: s.e. (D) Integrated septal HADA fluorescence increases similarly with labeling pulse duration in all strains. Error bars: s.e.

To test this hypothesis, we directly probed the spatial pattern of septal PG synthesis. We pulse-labeled WT cells and three mutants (E250A, D158A, and D212G) using the fluorescent D-alanine analog HADA (31). HADA is incorporated into the *E. coli* cell wall and can be used as a marker for nascent septal PG synthesis (31, 32). We reasoned that if treadmilling serves to time-average the spatial pattern of septum synthesis, labeling pulses shorter than or similar to the treadmilling period (~100 s) should result in punctate, incomplete septal labeling, rather than low level but homogenous labeling, whereas longer pulses would completely label the septum because of treadmilling. Indeed, we observed that for short labeling pulses, >60% and >80% of WT and D212G cells, respectively, displayed punctate incorporation of HADA at septa (Fig. 3B, C). With longer labeling pulses, the percentage of cells with incompletely labeled septa in WT cells and the mild GTPase mutant E250A (47% of WT GTPase activity and 63% of WT treadmilling speed) rapidly dropped for labeling pulses of ~100 s or longer (Fig. 3B, C). However, large fractions of cells of the more drastic mutants D158A (16% of WT GTPase activity, 27% of WT treadmilling speed) and D212G (14%, 6.7% respectively) still had incompletely labeled septa even after 1800 s (Fig. 3B, C). Interestingly, the total integrated fluorescence intensity of HADA at septa increased with longer labeling times, but there was no statistically significant difference between WT and the three mutants (Fig. 3D). While it remains to be verified that the incorporation rate of HADA is quantitatively proportional to the overall septal PG synthesis rate, we found that blocking the activity of either PBP1b or FtsI abolishes septal HADA incorporation (fig. S11), suggesting that HADA incorporation is specific to septal PG synthesis. Therefore, our labeling results are most consistent with the total septal PG synthesis activity not being significantly affected in FtsZ GTPase mutants, and instead the spatiotemporal distribution of synthesis is altered.

Next, we reasoned that since FtsZ recruits and scaffolds many proteins involved in septal synthesis, the spatiotemporal dynamics of the essential, septum-specific transpeptidase FtsI would likely follow that of FtsZ. We constructed a complementing (fig. S12), N-terminal fusion of FtsI to the photo-stable fluorescent protein TagRFP-T (33), and monitored its dynamics at visible constriction sites in FtsZ^WT^ and three FtsZ^mut^ backgrounds (D250A, D158A, and D212G) using wide-field epifluorescence microscopy. Strikingly, kymographs of TagRFP-T-FtsI fluorescence in FtsZ^WT^ cells showed clear diagonal tracks similar to that of FtsZ (Fig. 4A), indicating that FtsI exhibits processive redistribution along the septum. Most interestingly, in contrast to stationary FtsZ molecules in treadmilling FtsZ polymers (24), we found that individual TagRFP-T-FtsI molecules underwent directional movement along the septum (Fig. 4A, Movie S13). The movement was not unidirectional and exhibited large variations in time and in different cells (Fig. 4A, fig. S13, Movies S14-16). The mean speed of TagRFP-T-FtsI in FtsZ^WT^ cells was 19.3 ± 21.2 nm/s (µ ± s.d., *n* = 154 traces, Fig. 4E). In the three FtsZ^Mut^ strains, TagRFP-T-FtsI moved more slowly (Fig. 4B, C and D) with speeds that were linearly correlated with those of FtsZ treadmilling (*R*_Pearson_ = 1, p < 10^−45^, Fig. E). Thus, these data strongly indicate that FtsZ uses treadmilling powered by GTP hydrolysis to guide the directional movement of FtsI, and thereby direct the spatiotemporal distribution of septal cell-wall synthesis.

**Figure 4:**
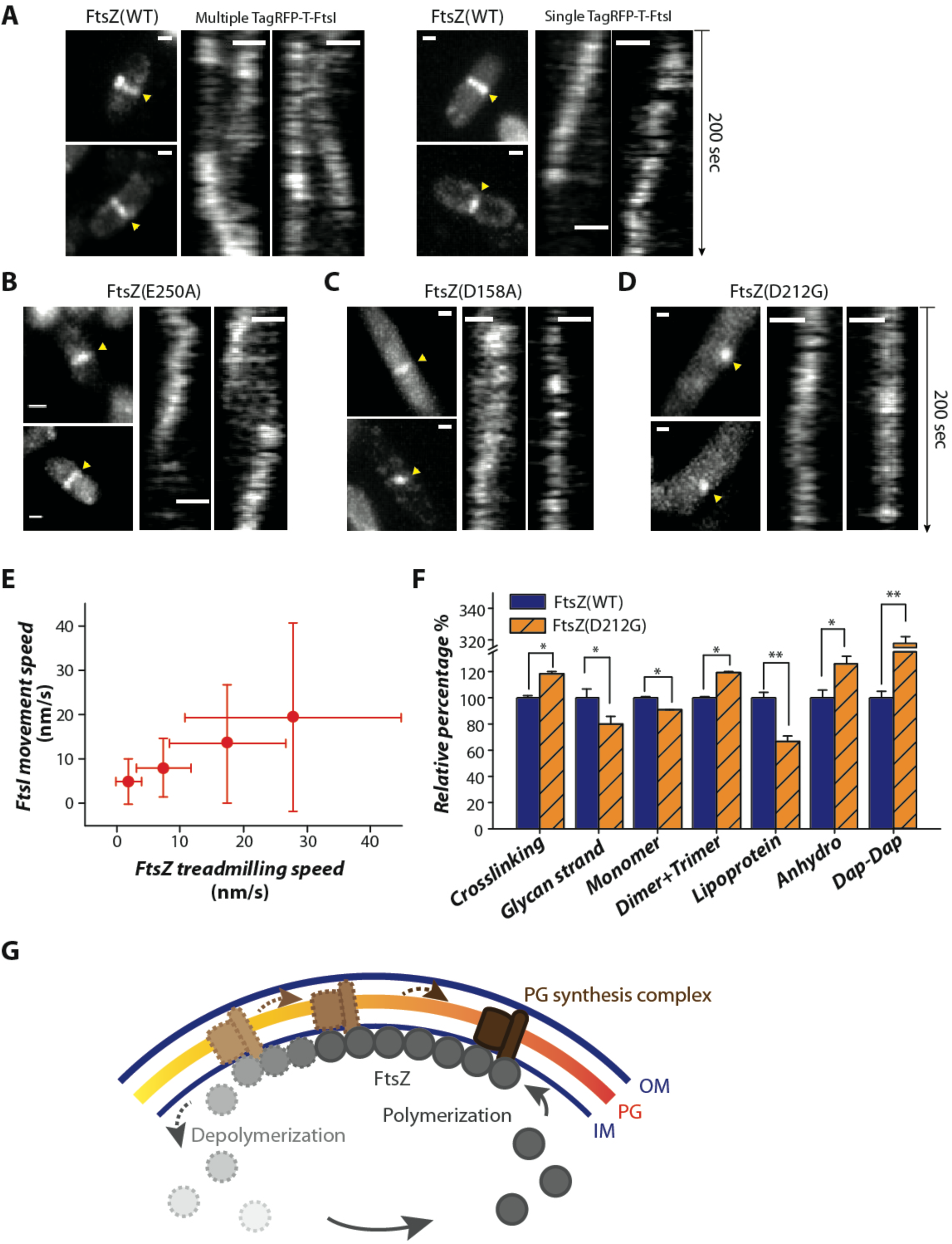
Altered directional movement of FtsI and septal PG composition in FtsZ^mut^ cells. (A) Directional movement of multiple (left) or single (right) TagRFP-T-FstI molecules along the septum in FtsZ^WT^ cells (see Movies S13 to S16). Maximum intensity projections of the whole cell are shown on the left, and the corresponding kymographs at positions denoted by arrows positions are shown on the right. (B-D) Directional movement of TagRFP-T-FtsI in FtsZ D250A (B), D158A (C) and D212G (D) strains. Note the punctate maximum intensity projection of TagRFP-T-FtsI (left) and the gradually decreasing movement speed (slope of the diagonal line) on the kymographs (right) in the order of D250A, D158A, and D212G strains. (E) The mean movement speed of TagRFP-T-FtsI in the four strains is highly correlated with FtsZ treadmilling speed (*R*_Pearson_ = 1, p = 10^−45^, Table S1). The slightly slower TagRFP-T-FtsI speed compared to that of FtsZ is due to the projection of a curved cell surface in the epi-fluorescence imaging mode of FtsI (12). (F) UPLC analysis of PG composition of WT (dark blue bars) and D212G (orange hatched bars) grown in M9 media. The relative percentage of each component of D212G was normalized to that of WT cells. Error bars: s.d. n=3. \**p* < 0.05, \*\**p* < 0.01 using unpaired t-test. (G) A schematic model depicting that the treadmilling of FtsZ polymers (dark and light gray circles) drives the directional movement (dashed arrows) of the septal PG synthesis machinery (brown rectangles), and hence leads to processive septal PG synthesis (yellow to red gradient indicates old to new PG).

Finally, since the spatial distribution pattern of cell-wall synthesis, the speed of FstI’s directional movement along the septum, and the resulting septal morphology were disrupted in FtsZ^mut^ cells, we investigated whether the biochemical composition and ultrastructure of the PG itself were altered. Using ultra performance liquid chromatography (UPLC), we quantified the proportion of each muropeptide species comprising cell walls isolated from WT and D212G cells, after growth in LB and minimal media (12). In both media, we observed that D212G cells have shorter glycan strands and higher crosslinking compared with WT (Fig. 4F, fig. S14A), indicating imbalances in the relative levels of glycan strand polymerization and crosslinking within the septal PG synthesis machinery of FtsZ^D212G^ cells. We also observed a large (~3-fold) increase in alternative Dap-Dap crosslinks (Fig. 4F), indicating that both the magnitude and the nature of crosslinking reactions was perturbed. To verify whether the difference could be due to the fact that D212G cells often have polarly localized Z-rings that produce round, DNA-less minicells, the PG composition of which essentially represents that of a cell pole, we compared the PG composition of minicells isolated from D212G and a BW25113 ∆*minC* strain; the latter also produces minicells due to misplacement of the Z-ring. We observed similar differences in PG composition in the minicells of both strains as with intact cells, indicating that the differences are not due to aberrant division site placement (fig. S14B). Interestingly, a *Caulobacter crescentus* FtsZ mutant lacking its C-terminal linker showed opposite changes (lower crosslinking and longer glycan strands), along with a septal cell wall bulging phenotype (34). Thus, the balance between glycan strand polymerization and crosslinking activities is likely important to define the shape of the septum (which eventually becomes the cell poles), and FtsZ likely coordinates the two enzymatic activities within the septal PG synthesis machinery.

Here, we have shown that FtsZ engages in treadmilling powered by GTP hydrolysis and organizes the spatial distribution and enzymatic activities of the septal PG synthesis machinery (a schematic model is shown in Fig. 4G). The broad conceptual similarities between FtsZ and MreB, with both illustrating complex dynamics of cytoskeletal proteins connected with cell-wall synthesis, suggest that coupling cytoskeletal motion to wall synthesis may be a general strategy across the kingdoms of life; indeed, movement of cellulose synthase complexes along cortical microtubules in plants mirrors the behavior of MreB and PG synthesis (35). However, the direction of causality for MreB and FtsZ are highly distinct: while MreB relies on wall synthesis for its movement, FtsZ exploits its innate treadmilling capacity to control the movement of septal synthesis. Treadmilling ensures evenly distributed wall synthesis through spatiotemporal averaging, loss of which can lead to aberrantly shaped or incomplete septa. Notably, a similar role has recently been demonstrated for the highly dynamic actomyosin contractile ring in fission yeast (36), suggesting a general connection between rapid reorganization of the division machinery and the need for robust polar morphogenesis. It remains to be discovered at the molecular level how FtsZ’s GTPase activity is coupled to treadmilling, how treadmilling directs FtsI movement, and whether other division proteins follow similar dynamics. In addition, it will be fascinating to probe whether the behaviors we have revealed are fully conserved across the bacterial kingdom.

## Acknowledgements

The authors thank lab members in the Xiao and Huang labs for valuable discussions and technical assistance, Erin Goley, Petra Levin, Carla Coltharp, Ganhui Lan, Ethan Garner and Georgia Squyres for critical discussions of the work, Erkin Kuru, Yves Brun, and Mike VanNieuwenhze for the HADA dye and assistance in its use, Joe Lutkenhaus for the MC123 strain, Harold Erickson for the FtsZ antibody, Erin Goley, Petra Levin, and their lab members for helpful advices on protein purification and GTPase assay, Roger Tsien for the TagRFP-T construct, and Michael Delannoy for assistance with SEM. This work was supported by NIH Director’s New Innovator Award DP2OD006466 (to K.C.H.), NSF CAREER Award MCB-1149328 (to K.C.H.), a National Science Foundation Graduate Student Fellowship (to A.M.), an Achievement Rewards for College Scientists Fellowship (to A.M.), NIH R01 GM086447 (to J.X.), National Science Foundation Grant EAGER MCB1019000 (to J.X.), and a Hamilton Smith Innovative Research Award (to J.X.). This work was also supported in part by the National Science Foundation under Grant PHYS-1066293 and the hospitality of the Aspen Center for Physics.

## References

1. E. Nogales, K. H. Downing, L. A. Amos, J. Lowe, Tubulin and FtsZ form a distinct family of GTPases. Nat Struct Biol 5, 451–458 (1998).

2. S. Vaughan, B. Wickstead, K. Gull, S. G. Addinall, Molecular evolution of FtsZ protein sequences encoded within the genomes of archaea, bacteria, and eukaryota. J Mol Evol 58, 19–29 (2004).

3. D. P. Haeusser, W. Margolin, Splitsville: structural and functional insights into the dynamic bacterial Z ring. Nat Rev Microbiol 14, 305–319 (2016).

4. J. Stricker, P. Maddox, E. D. Salmon, H. P. Erickson, Rapid assembly dynamics of the Escherichia coli FtsZ-ring demonstrated by fluorescence recovery after photobleaching. Proc Natl Acad Sci U S A 99, 3171–3175 (2002).

5. Y. Chen, K. Bjornson, S. D. Redick, H. P. Erickson, A rapid fluorescence assay for FtsZ assembly indicates cooperative assembly with a dimer nucleus. Biophys J 88, 505–514 (2005).

6. H. P. Erickson, Modeling the physics of FtsZ assembly and force generation. Proc Natl Acad Sci U S A 106, 9238–9243 (2009).

7. C. Coltharp, J. Buss, T. M. Plumer, J. Xiao, Defining the rate-limiting processes of bacterial cytokinesis. Proc Natl Acad Sci U S A 113, E1044–1053 (2016).

8. P. de Boer, R. Crossley, L. Rothfield, The essential bacterial cell-division protein FtsZ is a GTPase. Nature 359, 254–256 (1992).

9. D. RayChaudhuri, J. T. Park, Escherichia coli cell-division gene ftsZ encodes a novel GTP-binding protein. Nature 359, 251–254 (1992).

10. J. Stricker, H. P. Erickson, In vivo characterization of Escherichia coli ftsZ mutants: effects on Z-ring structure and function. J Bacteriol 185, 4796–4805 (2003).

11. Z. Lyu, C. Coltharp, X. Yang, J. Xiao, Influence of FtsZ GTPase activity and concentration on nanoscale Z-ring structure in vivo revealed by three-dimensional Superresolution imaging. Biopolymers 105, 725–734 (2016).

12. See supporting material online.

13. J. Buss et al., A multi-layered protein network stabilizes the Escherichia coli FtsZ-ring and modulates constriction dynamics. PLoS Genet 11, e1005128 (2015).

14. T. S. Ursell et al., Rod-like bacterial shape is maintained by feedback between cell curvature and cytoskeletal localization. Proc Natl Acad Sci U S A 111, E1025–1034 (2014).

15. S. van Teeffelen et al., The bacterial actin MreB rotates, and rotation depends on cell-wall assembly. Proc Natl Acad Sci U S A 108, 15822–15827 (2011).

16. E. C. Garner et al., Coupled, circumferential motions of the cell wall synthesis machinery and MreB filaments in B. subtilis. Science 333, 222–225 (2011).

17. J. Dominguez-Escobar et al., Processive movement of MreB-associated cell wall biosynthetic complexes in bacteria. Science 333, 225–228 (2011).

18. T. Baba et al., Construction of Escherichia coli K-12 in-frame, single-gene knockout mutants: the Keio collection. Molecular systems biology 2, 2006.0008 (2006).

19. C. Lu, J. Stricker, H. P. Erickson, Site-specific mutations of FtsZ-effects on GTPase and in vitro assembly. BMC Microbiol 1, 7 (2001).

20. H. A. Arjes, B. Lai, E. Emelue, A. Steinbach, P. A. Levin, Mutations in the bacterial cell division protein FtsZ highlight the role of GTP binding and longitudinal subunit interactions in assembly and function. BMC Microbiol 15, 209 (2015).

21. B. D. Bennett et al., Absolute metabolite concentrations and implied enzyme active site occupancy in Escherichia coli. Nature chemical biology 5, 593–599 (2009).

22. C. Lu, M. Reedy, H. P. Erickson, Straight and curved conformations of FtsZ are regulated by GTP hydrolysis. J Bacteriol 182, 164–170 (2000).

23. J. Buss et al., In vivo organization of the FtsZ-ring by ZapA and ZapB revealed by quantitative super-resolution microscopy. Mol Microbiol 89, 1099–1120 (2013).

24. M. Loose, T. J. Mitchison, The bacterial cell division proteins FtsA and FtsZ self-organize into dynamic cytoskeletal patterns. Nat Cell Biol 16, 38–46 (2014).

25. L. Niu, J. Yu, Investigating intracellular dynamics of FtsZ cytoskeleton with photoactivation single-molecule tracking. Biophys J 95, 2009–2016 (2008).

26. S. G. Addinall, J. Lutkenhaus, FtsZ-spirals and -arcs determine the shape of the invaginating septa in some mutants of Escherichia coli. Mol Microbiol 22, 231–237 (1996).

27. G. Fu et al., In vivo structure of the E.coli FtsZ-ring revealed by photoactivated localization microscopy (PALM). PLoS One 5, e12682 (2010).

28. M. P. Strauss et al., 3D-SIM super resolution microscopy reveals a bead-like arrangement for FtsZ and the division machinery: implications for triggering cytokinesis. PLoS Biol 10, e1001389 (2012).

29. S. J. Holden et al., High throughput 3D super-resolution microscopy reveals Caulobacter crescentus in vivo Z-ring organization. Proc Natl Acad Sci U S A 111, 4566–4571 (2014).

30. M. Jacq et al., Remodeling of the Z-Ring Nanostructure during the Streptococcus pneumoniae Cell Cycle Revealed by Photoactivated Localization Microscopy. mBio 6, (2015).

31. E. Kuru et al., In Situ probing of newly synthesized peptidoglycan in live bacteria with fluorescent D-amino acids. Angew Chem Int Ed Engl 51, 12519–12523 (2012).

32. A. K. Fenton, K. Gerdes, Direct interaction of FtsZ and MreB is required for septum synthesis and cell division in Escherichia coli. EMBO J 32, 1953–1965 (2013).

33. N. C. Shaner et al., Improving the photostability of bright monomeric orange and red fluorescent proteins. Nature methods 5, 545–551 (2008).

34. K. Sundararajan et al., The bacterial tubulin FtsZ requires its intrinsically disordered linker to direct robust cell wall construction. Nat Commun 6, 7281 (2015).

35. A. R. Paredez, C. R. Somerville, D. W. Ehrhardt, Visualization of cellulose synthase demonstrates functional association with microtubules. Science 312, 1491–1495 (2006).

36. Z. Zhou et al., The contractile ring coordinates curvature-dependent septum assembly during fission yeast cytokinesis. Mol Biol Cell 26, 78–90 (2015).

